# Tissue heterogeneity is associated with phenotypic but not genomic diversity in *Wolbachia* endosymbionts

**DOI:** 10.1101/2024.06.20.599863

**Authors:** Romain Pigeault, Yann Dussert, Raphaël Jorge, Theo Ulve, Marie Panza, Maryline Raimond, Carine Delaunay, Willy Aucher, Thierry Berges, David Ogereau, Bouziane Moumen, Jean Peccoud, Richard Cordaux

## Abstract

The mechanisms underlying within-host diversification in parasite populations are poorly understood. Yet, phenotypic and genotypic variation in parasites can shape their evolutionary trajectories and have important epidemiological consequences. Our aim was to determine whether the constraints associated with infecting different host tissues lead to the emergence and coexistence of multiple parasite sub-populations with distinct phenotypes. We tested this hypothesis using the most widespread bacterial endosymbiont, *Wolbachia*. We injected *Wolbachia* bacteria isolated from three tissues of the common pill-bug (*Armadillidium vulgare*) into uninfected individuals and monitored the growth rate and virulence of each bacterial sub-population in the new hosts. Our results highlight that within-host tissue heterogeneity leads to diverse *Wolbachia* phenotypes. High-depth genome re-resequencing of *Wolbachia* sub-populations revealed that this polymorphism was not due to genomic variation but was more likely a result of phenotypic plasticity. Indeed, we found no recurrent tissue-specific genomic variation among infected individuals. Our single nucleotide polymorphism (SNP) filtration pipeline, developed to ensure SNP validity, detected only one substitution. This *Wolbachia* variant, observed in one female, was present in all three bacterial sub-populations, with frequencies ranging from 24% to 58% depending on the tissue. Overall, our results support the stability of the *Wolbachia* genome with respect to the rarity of point mutations, in agreement with reports from other symbiotic systems. From a methodological perspective, our study highlights the need for considerable caution when detecting variants in endosymbiont populations, as our conservative approach led us to exclude more than 99.5% of the initially called variants.

## 1. Introduction

Among the many factors that influence species richness and community structure, environmental heterogeneity is generally considered to be a particularly important driver (Ben-Hur & Kadmon, 2020; Stein et al., 2014; Stein & Kreft, 2015). Since heterogeneous environments offer a variety of ecological niches and refuges from adverse conditions, species richness and persistence should increase with greater spatial heterogeneity (Ben-Hur & Kadmon, 2020; Burnett et al., 1998; Stein et al., 2014; Yang et al., 2015). Environmental heterogeneity also promotes genetic and phenotypic diversity within species through diversifying selection (Houle et al., 2020; B.-H. Huang et al., 2017; Mcdonald & Ayala, 1974; Rainey & Travisano, 1998), which may underpin local adaptations (Y. Huang et al., 2016; Qian et al., 2024; Trevail et al., 2021; van Houte et al., 2021). However, most studies that investigated relationships between environmental heterogeneity and diversity have focused on free-living species (e.g., (Langenheder & Lindström, 2019; Stein & Kreft, 2015)). The relationship between environmental heterogeneity and both endoparasites species diversity and phenotypic variability remains underexplored.

For endoparasites, environmental heterogeneity can exist both within individual hosts and across different hosts. Variation among individuals within host population is known to shape parasite evolution (Johnson et al., 2016; Regoes et al., 2000; White et al., 2020), driving rapid diversification (Johnson et al., 2016; Kamiya et al., 2014). However, a single host individual is treated by most theoretical and empirical studies on endoparasite evolution as a homogeneous environment – a single population of target cells without any structure – neglecting heterogeneities in cell type, immune response, nutrient supply or microbiota composition. Yet, several parasites exploit a heterogeneous environment composed of different biological tissues and fluids which form a spatially structured fitness landscape for the parasite. Successful colonization of different microenvironments, with their contrasted constraints, may involve the expression or selection of different parasitic phenotypes leading to within-host differentiation into several sub-populations (Didelot et al., 2016). This phenotypic variability can reflect phenotypic plasticity (e.g., expression plasticity, (Wernegreen & Wheeler, 2009), selection of genetic variants initially present in the inoculum (Chrostek & Teixeira, 2018) or of new variants arising from spontaneous mutations (Ailloud et al., 2019).

The co-occurrence of polymorphic sub-populations of parasites in different tissues or cell types has been described in several biological models. RNA viruses (e.g., plum pox virus, influenza, SARS-CoV-2, poliovirus) may display genetic diversity within single infected hosts, and co-existing viral variants may evolve differently in distinct cell types (Gupta et al., 2022; Jridi et al., 2006; Xiao et al., 2017). The ability of some RNA viruses to adapt to tissue-specific innate immune microenvironments may actually be critical for establishing robust infections (Xiao et al., 2017). Gastric biopsies from multiple stomach regions of *Helicobacter pylori*-infected hosts showed location-specific evolution of the bacteria, suggesting the existence of structured niches with distinct selective pressures within the stomach (Ailloud et al., 2019). In other cases, such as *Pseudomonas aeruginosa* or *Mycobacterium tuberculosis* chronic infections, the emergence of polymorphic sub-populations seems to be driven by tissue-specific adaptation (Jorth et al., 2015; Lieberman et al., 2016). Some within-host parasite sub-populations may constitute a reservoir from which new virulent variants can emerge (Bessière & Volmer, 2021), and which can delay clearance of parasite infection by the host immune system or drugs (Avettand-Fenoel et al., 2011; Clement et al., 2005; Obaldia et al., 2018). Understanding the evolutionary forces and mechanisms at the origin of within-host parasite diversity is essential, yet it is rarely studied. Descriptive studies (e.g., long-term ambulatory monitoring of chronically infected patients) cannot trace the divergence of variants found within a host and cannot identify the drivers of their diversification without controlling or assessing the composition of the infectious inoculum and ensuring the absence of subsequent infections. For this purpose, vertically transmitted parasites are particularly useful, as their transmission mode ensures the common origin of all within-host parasite sub-populations.

Among the most widespread vertically transmitted endosymbionts in animals is *Wolbachia pipientis*. This primarily maternally transmitted intracellular alpha-proteobacterium is widely distributed among arthropods and has been extensively studied with respect to its peculiar effects on its hosts (Kaur et al., 2021; LePage & Bordenstein, 2013). *Wolbachia* manipulates host reproduction in various ways, including the feminization of chromosomal male embryos, parthenogenesis, male killing, and sperm–egg incompatibility (Werren et al., 2008)). This endosymbiont can also have positive or negative effects on other aspects of the host’s life cycle (e.g., immunity, senescence) and influence the evolution of the host genome through horizontal gene transfer (Cordaux & Gilbert, 2017; Depeux et al., 2024; Sicard et al., 2014). The diversity of effects of *Wolbachia* on its hosts certainly relates to its wide distribution within infected individuals (Sicard et al., 2014). In addition to colonizing the female germline, ensuring vertical transmission, *Wolbachia* has indeed been observed in all major tissues (e.g., nerve chain, immune and digestive compartments, (Pietri et al., 2016)) of many arthropod species. The presence of vertically transmitted endosymbionts in somatic tissues may seem paradoxical, especially considering that systemic infection can decrease host fitness. Given this disadvantage (i.e., fitness alignment), these extensive somatic localizations must somehow benefit the parasite. Localization of *Wolbachia* in somatic tissues could influence host biology in such a way that favours its vertical and, possibly, horizontal transmission. However, the ability of the bacteria to colonize and establish in different tissular microenvironments could involve the expression or selection of different phenotypes.

To investigate this hypothesis, we assessed the relationship between tissue environment heterogeneity and phenotype variability in the *w*VulC *Wolbachia* strain, which infects the common pill-bug *Armadillidium vulgare* (Cordaux et al., 2004). We focused on three naturally infected tissues, which we selected for their physiological characteristics and the benefits that *Wolbachia* could gain by colonizing them: (i) gonads, which are a prime target for vertical transmission, (ii) haemocytes, as these circulating immune cells could contribute to spread *Wolbachia* to different host tissues and may facilitate its horizontal transmission (Braquart-Varnier et al., 2015), and (iii) the nerve chain, as the localization of *Wolbachia* in the brain and nerve chord could induce adaptive modifications of host behaviour (Bi & Wang, 2020; Templé & Richard, 2015). We hypothesized that the main phenotypic variation between bacteria sub-populations colonizing these tissues pertained to their replication rate. A high replication rate in vital tissues, such as the nerve chain, could disrupt tissue structure and reduce host lifespan (Le Clec’h et al., 2012; Strunov & Kiseleva, 2016). Thus, we expected *Wolbachia* populations from the host nerve system to show lower replication rates than those from the other tissues. We tested this hypothesis by injecting *Wolbachia* extracted from the three tissue types into uninfected *A. vulgare* individuals, and by monitoring their growth rates in their new hosts. We then tested whether *Wolbachia* sub-populations were adapted and not merely acclimated to their tissue types. In this case, we expected to find recurrent tissue-specific genomic variation among infected individuals. We tested this prediction by sequencing the whole genomes of focal bacterial sub-populations.

## 2. Materials and Methods

### 2.1 Biological models

All *A. vulgare* individuals used in this experiment were reared as previously described (Sicard et al., 2010). Given that the *Wolbachia* strain used in this study is feminizing, all the individuals used were females. As *Wolbachia* is not cultivable, phenotyping sub-populations requires trans-infection of an uninfected host with bacterial isolates from naturally infected animals. For this purpose, we used a common pill-bug source line (WXw) infected by the *w*VulC *Wolbachia* strain (source individuals) and an uninfected recipient line (WXa, recipient individuals) into which we have injected bacteria. The two laboratory lines came from the same original population sampled in Helsingør (Denmark).

### 2.2 Experimental infection

To characterize the phenotype of the different *Wolbachia* sub-populations associated to three different tissues of their native host, five batches of trans-infection were carried out. Three batches (i.e., three biological replicates) were used to monitor early infection dynamics within the recipient host (day 0 to day 60 post-infection, trans-infection experiment 1), and two batches were used to estimate bacterial persistence in tissues (i.e., two biological replicates, day 200 post-infection, trans-infection experiment 2). For each batch, one tissue suspension was prepared from each of the following tissues: haemolymph, ovaries and nerve chain. Tissues were collected from five source females from the WXw line (infection status previously validated by quantitative PCR, see section 2.4). For each suspension, tissues were crushed in 1 mL of Ringer solution (1.4 mM CaCl2, 2.4 mM HNaCO3, 2 mM KCl, 0.4 M NaCl) for haemolymph and 2 mL for ovaries and nerve chain. The resulting suspension was filtered through a 1.2 µm pore membrane to remove cell debris and host cell nuclei. A prior experiment conducted on 15 females from the source line (WXw) allowed us to highlight that the number of bacteria per ng of DNA differed between the three target tissues (see **Fig. S1** in appendix 1). Therefore, the filtered solutions were diluted in Ringer solution to adjust the final concentration of bacteria at 2500 ± 300 cells/µL for the trans-infection batches used to describe *Wolbachia* infection dynamics, and at 1282 ± 205 cells/µL for the two batches made to investigate bacterial persistence 200 days after injection. The variation in *Wolbachia* concentration between the two experiments stems from their differing timing, with the first conducted in January and the second in April. Since *Wolbachia* concentration (number/ng of DNA) can fluctuate based on host size and, consequently, age, this resulted in slight differences in *Wolbachia* concentration between the two trans-infection experiments. Another prior experiment showed that the proportion of live bacteria in the different filtered tissue solutions was similar (LRT = 4.0279, p = 0.1335, proportion of live *Wolbachia* ± 95%CI, haemolymph = 0.815 ± 0.017, nervous chain = 0.875 ± 0.014, ovaries = 0.862 ± 0.037, see appendix 1 for a detailed protocol).

One µL of each filtrate was injected using a thin glass needle into the general cavity of each recipient individual through a small hole pierced at its posterior part. Each tissue filtrate from the three trans-infection batches used to study *Wolbachia* infection dynamics (trans-infection experiment 1) was injected into 21 recipient females of the WXa line, resulting in a total of 63 females transinfected per tissue filtrate. Then, on days 20, 40 and 60 post-injection, 6-7 recipient females per trans-infection batch were randomly selected to quantify *Wolbachia* in the focal tissues (see section 2.4). To study bacterial persistence in recipient host tissues (trans-infection experiment 2), 10 females were injected with each tissue filtrate for each of the two trans-infection batches. Two hundred days post-injection, all surviving females were dissected to quantify *Wolbachia* in the three focal tissues (see section 2.4). For each trans-infection batch, control groups were created using the same protocol, but with female sources originating from the WXa line.

### 2.3 Mortality monitoring

The effect of *Wolbachia* injection on pill-bug mortality was monitored every 10 days as part of the study of the influence of *Wolbachia* tissue origin on the dynamics of early infection within the recipient host (trans-infection experiment 1), and every 15-25 days as part of the study of *Wolbachia* tissue origin on the persistence of the bacterium in tissues (trans-infection experiment 2). When individuals were harvested during the experiment to quantify *Wolbachia*, they were censored for the survival analysis (see section 2.9).

### 2.4 Quantification of *Wolbachia* in source and recipient host’s tissues

To compare *Wolbachia* density in different tissues of source line and recipient hosts, total DNA was extracted from the haemolymph, nerve chain and ovaries of individuals using standard protocols (Qiagen DNeasy 96 Blood & Tissue kit). For each sample, the purity of the extracted DNA (OD 260/280 nm and 260/230 nm ratios) was measured using a Nanodrop 1000 spectrophotometer (Thermofisher). To quantify *Wolbachia* load, we developed a fluorescent probe-based quantitative PCR (qPCR) approach amplifying a *Wolbachia* locus and a reference host nuclear locus in the same reaction. We used primers wsp208f (5’- TGG-TGC-AGC-ATT- TAC-TCC-AG-3’) and wsp413r (5’-TCG-CTT-GAT-AAG-CAA-AAC-CA-3’) targeting the *Wolbachia* protein surface gene (*wsp*, (W. L. Le Clec’h et al., 2012)) and we designed primers amplifying a portion of the single-copy nuclear gene encoding the mitochondrial leucine-tRNA ligase of *vulgare*: TLeuF (5’- TGT-ACA-CAT-CGA-GCA-GCA-AG-3’) and TLeuR (5’- AAA-GAG-GAG-CGG-AGA-GTT-TCA-G-3’) (Durand et al., 2023). The *Wolbachia wsp* double-dye probe (5’-TTG-CAG- ACA-GTG-TGA-CAG-CGT-T-3’) was labelled with Hexachlorofluorescein (HEX) as a reporter at the 5’ end and Black Hole Quencher 1 (BHQ-1) at the 3’ end. *A. vulgare* TLeu probe (5’-ACG- AAG-TTC-GCC-CTG-TTC-TGG-A-3’) was labelled with 6-carboxy-fluorescein (FAM) as a reporter and BHQ-1 as a quencher. The qPCR reactions were performed using Roche LIGHTCYCLER 480 with the following program: 2 min at 50°C, 10 min at 95°C and 45 cycles, 15 s at 95°C, and 1 min at 60°C. For each sample, two independent technical replicates were carried out and the mean cycle-threshold value was calculated. *Wolbachia* load was then calculated relative to the reference gene (TLeu) using the delate CP method: 2^-(CP(*wsp*) – CP(TLeu))^.

### 2.5 Genome assembly of *Wolbachia w*VulC

To study the within-host genetic diversity of *Wolbachia*, we first assembled the genome of the *w*VulC *Wolbachia* strain from the WXw line. Three female individuals were used for Oxford Nanopore Technologies (ONT) long-read sequencing and one for Illumina short-read sequencing. Total DNA from haemolymph, nerve chain and ovaries of each female was extracted using standard protocols (DNA used for ONT sequencing: Macherey-Nagel NucleoBond HMW DNA kit, DNA used for Illumina sequencing: Qiagen DNeasy 96 Blood & Tissue kit). The long-read sequencing library was prepared using the SQK-LSK109 Ligation Sequencing Kit (ONT) and sequenced with an R9.4.1 flow cell on a MinION Mk1B (ONT). Reads were basecalled with Guppy 6.3.8 (RRID:SCR_023196) using the SUP model. The short-read library was prepared using the TruSeq Nano DNA Library Prep kit (Illumina) and sequenced on a HiSeq X (Illumina, paired-end 2 × 150 bp).

ONT adapters were first trimmed with porechop 0.2.4 (https://github.com/rrwick/Porechop), and reads were filtered using NanoFilt 2.7.1 (minimum average quality: 7, minimum length: 1000 bp) implemented in NanoPack (De Coster et al., 2018). Reads were then assembled with Flye 2.8.3 (Kolmogorov et al., 2019) using default parameters. The entire assembly was first polished with the ONT reads using medaka 1.7 (https://github.com/nanoporetech/medaka, r941_min_fast_g507 model, 2 rounds). For assembly polishing using short reads, Illumina reads were mapped onto the assembly with bwa-mem2 2.2.1 (Vasimuddin et al., 2019) and used for further polishing with Polypolish 0.4.3 (Wick & Holt, 2022). This step was carried out twice. The largest assembled contig was reported detected as circular by the Flye assembler and had the expected length (∼1.6 Mb) of the *w*VulC *Wolbachia* genome. We aligned this contig against a previous version of the genome (GenBank accession: ALWU00000000.1) with nucmer in MUMmer 4.0.0rc1 (Marçais et al., 2018), confirming that the contig corresponded to the *w*VulC strain. After mapping the short reads for a third time onto this contig, short variants were called with Freebayes 1.3.1 (Garrison & Marth, 2012). Variants were then visually reviewed using the IGV genome browser (version 2.14.0) (Robinson et al., 2017). Eight of them, corresponding to errors, were kept. We corrected the genome sequence with these eight variants using the consensus command in BCFtools 1.17 (Danecek et al., 2021). Following the recommendations of Ioannidis (Ioannidis et al., 2007), the genome sequence was rotated to start at the *hemE* gene using the fixstart command of Circlator 1.5.5 (Hunt et al., 2015). The genome was annotated with the NCBI Prokaryotic Genome Annotation Pipeline (PGAP) 2022-12-13.build6494 (Haft et al., 2024; Li et al., 2021; Tatusova et al., 2016). A search of Insertion Sequence (IS) elements was carried out with digIS v1.2 (Puterová & Martínek, 2021). Additionally, bacteriophage-derived regions were detected using PHASTEST (Wishart et al., 2023). Results of these annotations were plotted with Circos v0.69-9 (Krzywinski et al., 2009). The completeness of the genome and the gene annotation was assessed using BUSCO 5.3.2 (Manni et al., 2021) using the rickettsiales_odb10 dataset.

### 2.6 Whole-genome resequencing and variant calling

Three *A. vulgare* females from the WXw line were used to study the genetic diversity of the different *Wolbachia* sub-populations associated with the nerve chain, ovaries and haemolymph. Two of these females were sisters (names: F1_2015 & F2_2015). The matriline of the third one (F_1999) diverged from the matriline of the sisters for 18 generations (i.e., 18 years). Females were dissected and their tissues isolated individually. Total DNA of each biological sample was extracted using the Qiagen DNeasy 96 Blood & Tissue kit. DNA purity and quantity were measured as described above. The haemolymph of two females could not be sequenced because it did not yield enough DNA. DNA libraries were prepared using the TruSeq Nano DNA Library Prep kit and sequenced on the Illumina HiSeq 2000 platform to generate 2×150 bp reads, by Genoscreen (Lille, France). Read quality was assessed with FastQC 0.11.9 (http://www.bioinformatics.babraham.ac.uk/projects/fastqc). Fastp 0.23.0 (Chen et al., 2018) was used to remove adapter contamination and systematic base calling errors. To mitigate misalignments of reads originating from the host genome, notably from *Wolbachia*-derived nuclear insertions (Chebbi et al., 2019; Leclercq et al., 2016), reads were aligned using bwa-mem2 (Vasimuddin et al., 2019) against a reference including our *w*VulC genome assembly, the assembly of the partial mitochondrial genome of *A. vulgare* (GenBank accession number: MF187614.1, (Peccoud et al., 2017)) and the genome assembly of *A. vulgare* (accession number: GCA_004104545.1, (Chebbi et al., 2019)). PCR duplicates were marked with samtools markdup (v1.14). Read coverage along the *w*VulC genome was computed for each sample for adjacent 5 kb windows using mosdepth 0.2.9 (Pedersen & Quinlan, 2018). For a given sample, normalized coverage for each window was calculated by dividing the coverage value by the median of all values in R (v. 4.2.1) (R Core Team, 2022). SNPs and short indels were detected using Freebayes 1.3.1 (Garrison & Marth, 2012), with a filter on the minimum fraction of observations supporting a variant (F = 0.03), disabling population priors (-k argument) and using an aggregate probability of observation balance between alleles (-a argument). Variants were then filtered using a custom R script. This script selected only variable positions for which the read total depth was over 200 and for which the alternative allele was supported by at least 10 reads within a tissue (*Wolbachia* genome coverage - mean ± se: ovaries = 723.9 reads ± 134.8, nerve chain = 769.8 ± 81.0, haemolymph = 533.7).

### 2.7 Variant validation

As the *A. vulgare* lineage used in our experiments may contain nuclear inserts absent from the reference genome used for read mapping (Chebbi et al., 2019) the risk that sequences from *Wolbachia*-derived nuclear inserts generated spurious SNPs may not have been eliminated. To exclude remaining spurious SNPs, we implemented a three-step approach. (**1**) We checked that the sequences containing candidate SNPs were absent from uninfected individuals sequenced in previous studies, using all short-read data generated by (Chebbi et al., 2019) and (Cordaux et al., 2021). These data constitute whole genome sequences of six pools of 10 males or females and five individuals, from three uninfected lineages. We aligned these short reads with bwa-mem2 on the *w*VulC reference sequence. On the 11 resulting alignment files, variant calling was carried out with the same parameters as above, but only at positions of detected candidate *Wolbachia* variants. All variants for which the alternative allele was present in the reads sequenced from uninfected individuals were eliminated. We then defined a 300-bp window around the position of each remaining variant (starting 150 bp before the site) and excluded variants in windows containing at least two reads from uninfected individuals. (**2**) To further check that each remaining variable region was absent from uninfected individuals, we then designed PCR primers to amplify 120-250 bp targets encompassing the focal variant. These primers were tested on two females from the recipient lineage (i.e., uninfected by *Wolbachia*), on two females from the source line (i.e., infected by *Wolbachia*) and on two of their brothers (not infected by *Wolbachia,* since infection induces feminization). If the primers amplified DNA from at least one non-infected host, then the variant was eliminated. (**3**) Finally, we checked whether sequencing reads carrying non-reference SNP alleles represented sequences that were exceedingly divergent from the reference strain (e.g., insertions of *Wolbachia* DNA into the host genome, or a related bacterium). To do so, we automatically analyzed every candidate SNP via a script that parses the alignment (.bam) files of reads generated in this experiment. This script records the proportion of clipped reads, the proportion of read pairs that were aligned in proper pairs and the number of mismatches between reads and the reference (excluding the SNP itself). The script then performs Fisher exact tests to evaluate if these proportions significantly differ between reads carrying the reference allele and reads carrying the alternativeallele. Any SNP for which the p-value of any test was below 5% (after false discovery rate correction) was excluded. Lastly, the sequencing depth around SNPs that had passed all filters was averaged over 100-bp windows, via samtools depth, and plotted to check for anomalies, mainly the presence of collapsed repeated regions not fully resolved in the *Wolbachia* reference genome assembly.

To predict the effect of retained SNPs and indels on proteins coded by annotated genes, we used SNPeff 5.2c (Cingolani et al., 2012). Functional domains in these proteins were annotated using the InterProScan online service (http://www.ebi.ac.uk/interpro/, database release 100.0, (Paysan-Lafosse et al., 2023)).

### 2.8 Screening for variants in maternal lines

To determine whether the *Wolbachia* variant(s) validated by our filtration procedure (see above) arose during the focal individuals’ lifetime or were maternally inherited, we searched for these variants in their maternal lineage. Specifically, we extracted DNA from the ovaries of sisters and maternal ancestors of focal individuals, going back six generations thanks to our broodstock collection stored at -20°C. The variants as well as the *Wolbachia* reference lineage were searched by amplification-refractory mutation system (ARMS, (C. R. Newton et al., 1989)). For this purpose, for each variant we designed a couple of primers targeting either the reference allele or the variant allele (see primers in appendix 1). ARMS analyses were performed in a 25 µl reaction volume containing 0.5 µl genomic DNA, 5 µl 5X buffer (Promega), 0.5 µl dNTP, 1.25 µl forward and reverse primers and 0.125 µl GoTaq® polymerase (Promega). The volume was adjusted to 25 µl with double-distilled water. PCR regimen was as follows: initial denaturation at 95°C for 5 min, followed by 35 cycles for 30 s at 95°C, 30 s at 61°C, 30 s at 72°C, and then a final extension for 5 min at 72°C, and finally the PCR products were maintained at 4°C in the end. PCR products were separated on a 1.5% agarose gel (23min., 100V).

### 2.9 Statistical Analyses

Statistical analyses were carried out using the R software (R Core Team, 2022). The influence of the tissular origin of the injected *Wolbachia* on *A. vulgare* mortality was evaluated using the ‘survreg’ function with either exponential or Weibull errors distribution (‘survival’ package, (Therneau et al., 2024)). The model included survival as a response variable, and infection status (i.e., injected with *Wolbachia*-infected or uninfected tissue filtrate), tissue origin of the injected filtrate and experimental block as explanatory variables. The influence of *Wolbachia* tissue origin on its ability to colonize and infect different focal tissues of recipient hosts over time was assessed using a generalized linear mixed model (GLMM). Log-transformed relative quantification (RQ) values representing the per-host cell *Wolbachia* dose were fitted as a response variable, assuming normal error distribution. The tissular origin of *Wolbachia*, the focal tissue of recipient injected hosts and the day post injection were used as explanatory variables. The quadratic term day post injection was added to assess whether it significantly improved the model fit. Individual IDs nested in experimental block were specified as random-effect variables in the model. To compare *Wolbachia* loads measured in the tissues of recipient hosts six months after experimental infection with that measured in the same tissues from females of the same age but naturally infected by *Wolbachia* (from the WXw line), we ran, for each tissue, a linear model with log-transformed relative quantification (RQ) values fitted as a response variable and the origin of the individuals (i.e., naturally infected or experimentally infected) as an explanatory variable.

Maximal models including all higher-order interactions were simplified by sequentially eliminating non-significant terms and interactions to establish a minimal model (Crawley, 2012). The significance of the effects of explanatory variables was established using a likelihood ratio test (LRT, (Crawley, 2012)) whose statistics approximately follows a Chi-square distribution (Bolker, 2008) or an F test. The significant LRT or F value given in the Results section are for the minimal model, whereas non-significant values correspond to those obtained before the deletion of the variables from the model. *A posteriori* contrasts were carried out by aggregating factor levels together and by testing the fit of the simplified model using a likelihood-ratio test (Bolker, 2008; Crawley, 2012).

## 3. Results

### 3.1 *Wolbachia* infection does not significantly influence host survival

Under the assumption of both exponential errors and non-constant hazard, no effect of *Wolbachia* tissue origin on the survival rate of *A. vulgare* during early stage of infection (from day 0 to 60, trans-infection experiment 1) was detected (exponential errors: LRT = 0.600, p = 0.740, Weibull errors: LRT = 0.602, p = 0.741, **Fig. 1A**). Overall, the injection of *Wolbachia*, irrespective of its tissue origin, was not shown to influence the survival of individuals (exponential errors: LRT = 1.040, p = 0.308, Weibull errors: LRT = 1.039, p = 0.307). Sixty days after injection, the survival rate of infected individuals was 0.87 (± 95% CI, ± 0.79-0.96), compared to 0.80 (± 0.71-0.91) for control animals.

**Figure 1.**
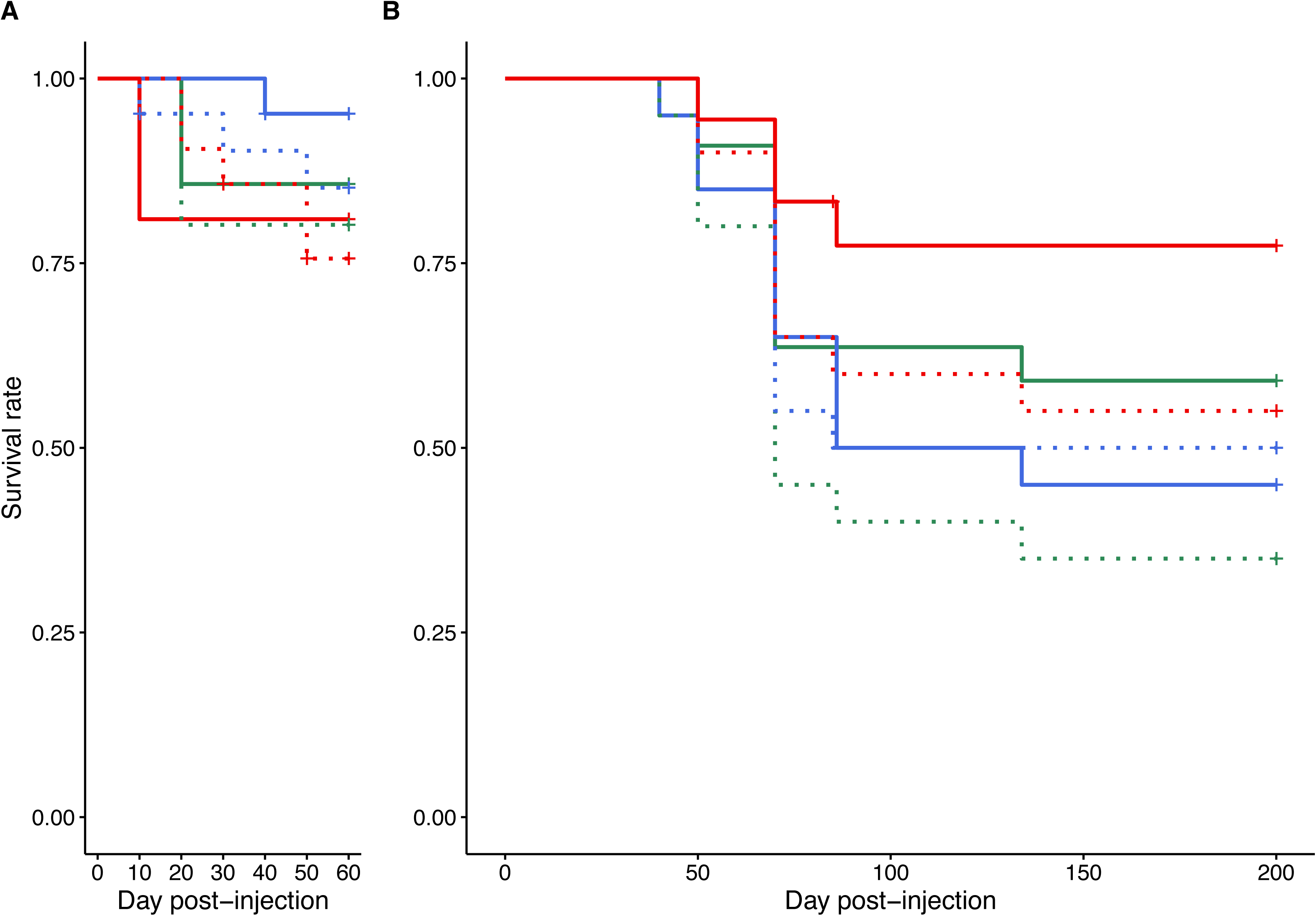
Survival rate of *Armadillidium vulgare* following *Wolbachia* (*w*VulC) trans-infection. Effect of tissue origin of injected *Wolbachia* on mortality of recipient hosts monitored (**A**) every 10 days from day 0 to day 60 post-injection and (**B**) every 15-20 days from day 0 to day 200 post-injection. Solid lines: recipient hosts injected with crushed filtered tissue from animals naturally infected by *Wolbachia w*Vulc. Dotted lines: control individuals injected with crushed filtered tissue *from A. vulgare* not infected by *Wolbachia.* The three colours correspond to the tissue of origin of Wolbachia. Green: nerve chain, red: ovaries, blue: haemolymph

In the trans-infection batches used to investigate the influence of *Wolbachia* tissue origin on its ability to persist in recipient host tissue (trans-infection experiment 2), the mortality of animals was also not significantly influenced by *Wolbachia* tissue origin (exponential errors: LRT = 2.65, p = 0.265, Weibull errors: LRT = 2.88, p = 0.236) or even just by the fact of being infected by the bacteria (whatever the origin of the tissue, exponential errors: LRT = 2.22, p = 0.136, Weibull errors: LRT = 2.45, p = 0.117, **Fig. 1B**). The survival rate of infected individuals 200 days after the experimental infection was 0.59 ± 0.48-0.74 and 0.47 ± 0.36-0.61 for infected and control animals respectively.

### 3.2 The tissue origin of *Wolbachia* influences its early infection dynamics in recipient host tissues (trans-infection experiment 1)

After injection, the presence of *Wolbachia* was detected by qPCR in the three tested tissues of all recipient hosts, confirming its ability to colonize a new host. *Wolbachia* density increased over time (post-injection day: LRT = 16.239, p < 0.0001) and fitting the quadratic term (day_post_injection²) strongly improved model fit (LRT = 32.181, p < 0.0001), suggesting that *Wolbachia* burden was more accurately modelled by an accelerated polynomial function of day post-injection (**Fig. 2A**). The density of *Wolbachia* in tissues started very low on day 20 post-injection, then rose slightly between day 20 and day 40, to drastically increase between day 40 and day 60 post-injection. Although the pattern of infection in the different recipient tissues was overall similar, *Wolbachia* load was significantly influenced by the interaction between the colonized recipient tissue and the day post-injection (LRT = 74.309, p < 0.0001). The dynamics of infection within the nerve chain were significantly different from those observed in the haemolymph and ovaries, leading to a higher bacterial load in nerve tissues 60 days post-infection (LRT = 57.228, p < 0.0001, LRT = 122.240, p < 0.0001, respectively, **Fig. 2A**). The dynamics of infection were also significantly different within haemolymph and ovaries (LRT = 17.771, p = 0.0013, **Fig. 2A**), as bacterial load was lower in ovaries at the end of monitoring (mean RQ ± se, nerve chain = 6.582 ± 1.596, haemolymph = 4.864 ± 1.612, ovaries = 2.533 ± 0.594, **Fig. 2A**). *Wolbachia* infection dynamics were also influenced by the tissue origin of the injected bacteria (interaction between days post-injection and tissue origin: LRT = 24.205, p < 0.0001). Whether the bacteria came from the haemolymph or ovaries of naturally infected animals did not significantly affect the colonization dynamics of *Wolbachia* in the injected recipients (LRT = 0.647, p = 0.723, **Fig. 2B**). However, bacteria from the nerve chain colonized recipient hosts less rapidly than bacteria from other tissues (nerve chain versus haemolymph: LRT = 28.981, p < 0.0001, nerve chain versus ovaries: LRT = 33.64, p < 0.0001). At 60 days post-injection, the *Wolbachia* load was on average more than twice as high when the bacteria came from the ovaries and haemolymph than when they came from the nerve chain of infected source hosts (mean RQ ± se, ovaries = 8.191 ± 2.18, haemolymph = 5.26 ± 1.077, nerve chain = 0.899 ± 0.197, **Fig. 2B**). However, there was no significant interaction between the colonized recipient tissue and the *Wolbachia’s* original tissue, suggesting that bacteria originating from a tissue do not colonize that same tissue more effectively in recipient hosts (LRT = 6.957, p = 0.138).

**Figure 2.**
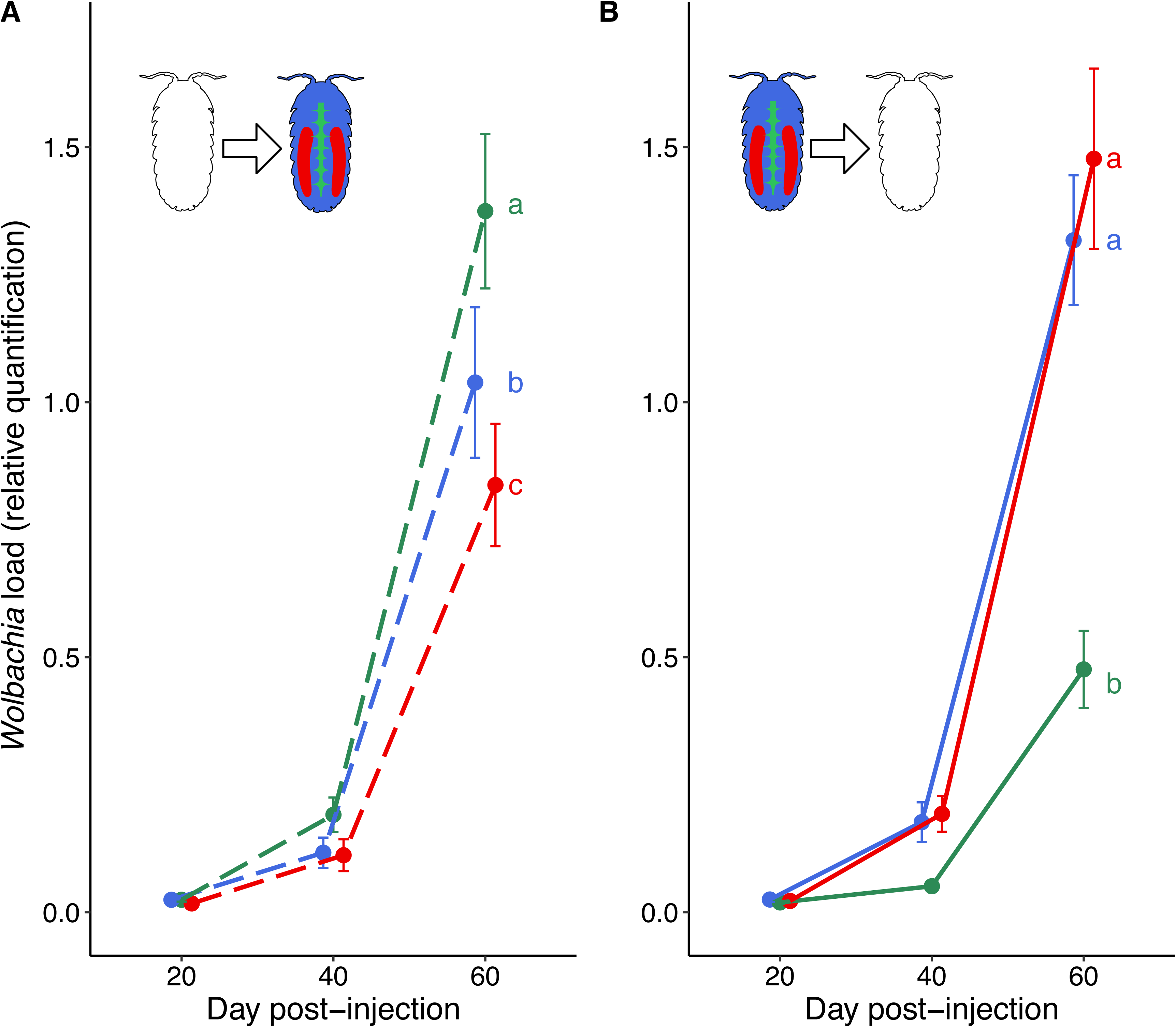
Infection dynamics of *Wolbachia* (*w*VulC) in *Armadillidium vulgare* from day 0 to day 60 post-injection. (**A**) *Wolbachia* loads in nerve chain (green), haemolymph (blue) and ovaries (red) of recipient hosts trans-infected by *Wolbachia*, all source tissue considered. (**B**) Influence of *Wolbachia* tissue origin on infection dynamics in tissue loads in recipient host, all recipient tissue considered. Green: nerve chain, red: ovaries, blue: haemolymph. Error bars correspond to the standard error. Infection dynamics not connected by the same letter are significantly different (p < 0.05).

### 3.3 The tissue origin of *Wolbachia* does not influence significantly its persistence in recipient host tissues (trans-infection experiment 2)

*Wolbachia* densities at 200 days after infection were significantly different between tissues from the recipient hosts (LRT = 39.224, p < 0.0001, **Fig. 3**). Bacterial load was the highest in the nerve chain and the lowest in the haemolymph (contrast analyses, haemolymph versus nerve chain: LRT = 38.133, p < 0.0001, haemolymph versus ovaries: LRT = 20.823, p < 0.0001, nerve chain versus ovaries: LRT = 6.969, p = 0.008, **Fig. 3**). The bacterial load in the ovaries and haemolymph of recipient females were similar to those observed in the same tissues of females naturally infected by *Wolbachia* (source lineage, F = 0.184, p = 0.672, F = 0.268, p = 0.609, respectively, **Fig. 3 & 4**). In contrast, bacterial density in the nerve chain of experimentally infected females was on average more than twice that measured in the nerve chain of naturally infected individuals (F = 38.089, p < 0.0001, **Fig. 3 & 4**).

**Figure 3.**
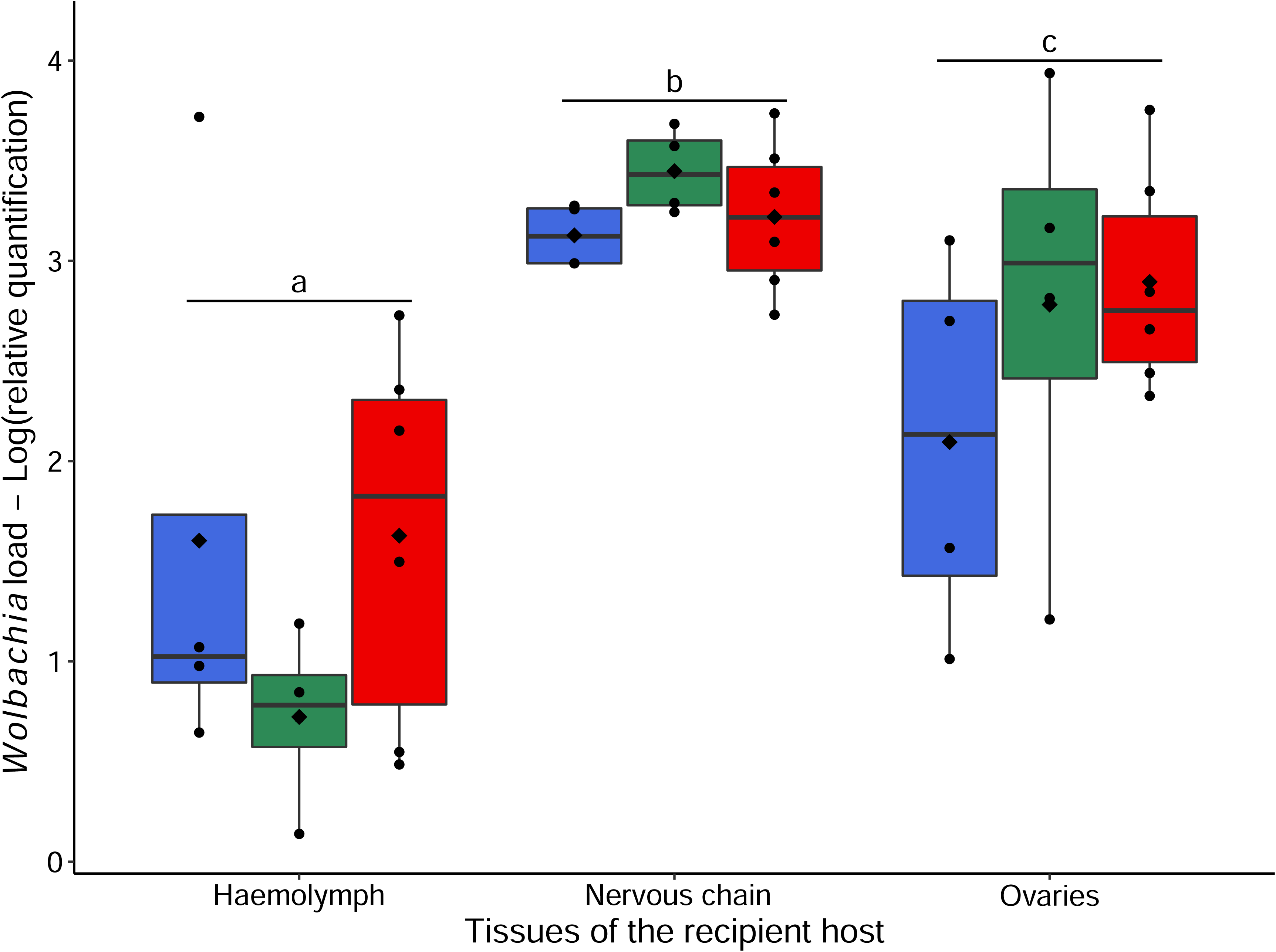
*Wolbachia* (*w*VulC) load in *Armadillidium vulgare* tissues six months post-injection. The coloured boxplots illustrate the tissue origin of Wolbachia (green: nerve chain, red: ovaries, blue: haemolymph). However, the analysis did not reveal any significant effect of the tissue origin of the bacteria on the intensity of infection (RQ) in the recipient host tissues. Boxes above and below the medians (horizontal lines) show the first and third quartiles, respectively. Black diamonds represent the means. Levels not connected by the same letter are significantly different (p < 0.05).

**Figure 4.**
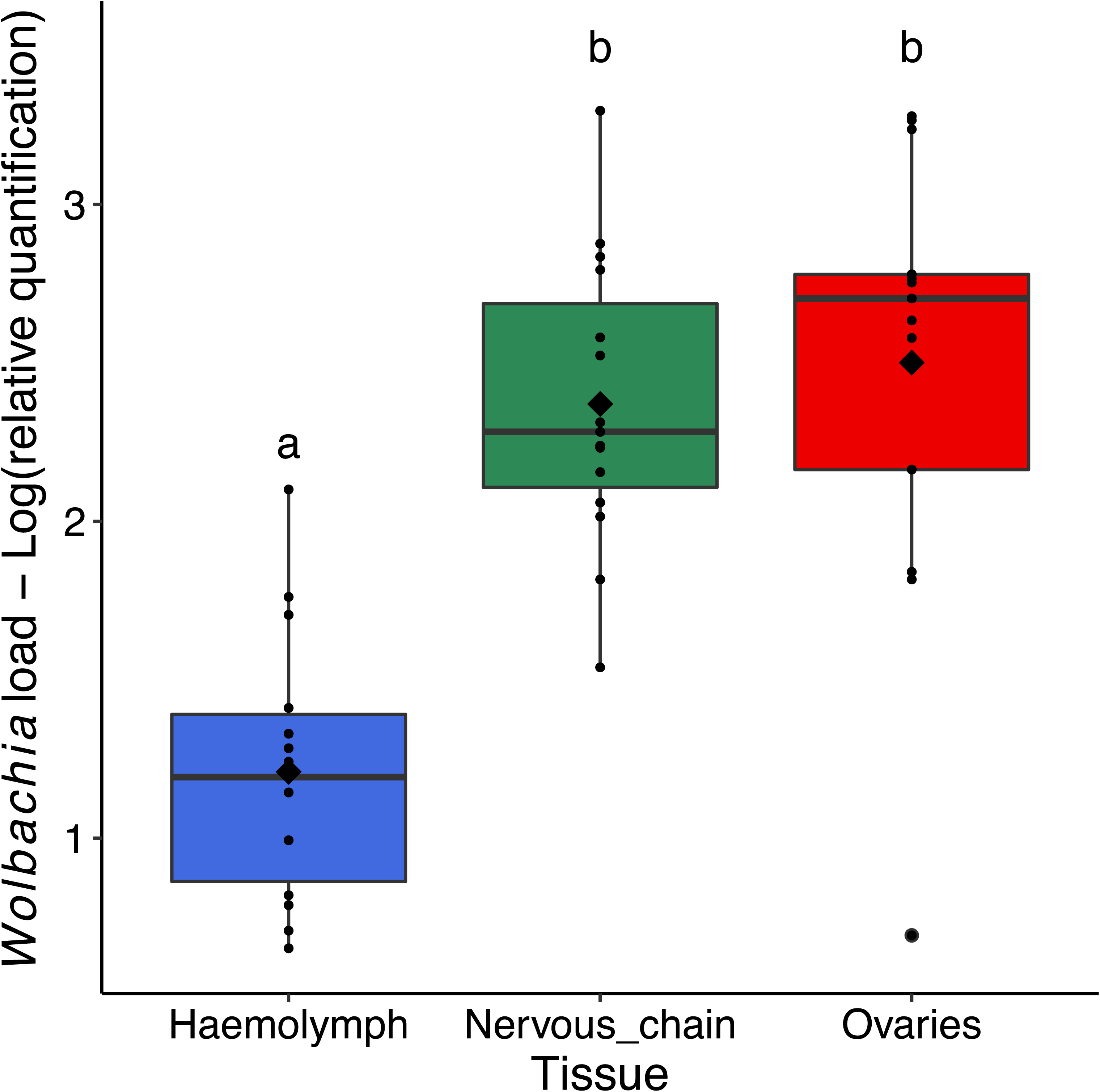
*Wolbachia* (*w*VulC) load in tissues from naturally infected females (source line). *Wolbachia* load was measured in the haemolymph, nerve chain and ovaries of 15 females from the source line. The analysis showed that the bacterial load was lower in the haemolymph than in the ovaries and nerve chain. Boxes above and below the medians (horizontal lines) show the first and third quartiles, respectively. Black diamonds represent the means. Levels not connected by the same letter are significantly different (p < 0.05). Green: nerve chain, red: ovaries, blue: haemolymph.

Contrary to what was observed during the early phase of infection, there was no effect of the tissue origin of *Wolbachia* on the bacterial load in recipient tissues at 200 days post-injection (LRT = 0.368, p = 0.832). Similarly, *Wolbachia* density in recipient tissues was not influenced by the interaction between the colonized tissue and the tissue origin of the injected bacteria (LRT = 9.539, p = 0.059).

### 3.4 Complete genome assembly of the *w*VulC *Wolbachia* strain

Using ONT reads, a complete circular genome sequence (1,638,144 bp) of the *w*VulC *Wolbachia* strain was assembled. The estimated completeness of the assembly was very high, with 99.8% of the expected BUSCO genes found in the sequence (C: 99.8% [S: 99.5%, D: 0.3%], F: 0.0%). The annotation of the genome resulted in 1,390 protein-coding sequences and 208 pseudogenes (**Fig. 5**). The *w*VulC genome contained 138 IS elements that are larger than 150 bp, representing 8.45% of the total length of the genome. This high proportion of repeated sequences for a bacterium is similar to that of other *Wolbachia* strains (Cerveau et al., 2011). We also identified seven prophage regions (six intact and one questionable region, following PHASTEST scoring criteria), representing 12.1% of the *w*VulC genome.

**Figure 5.**
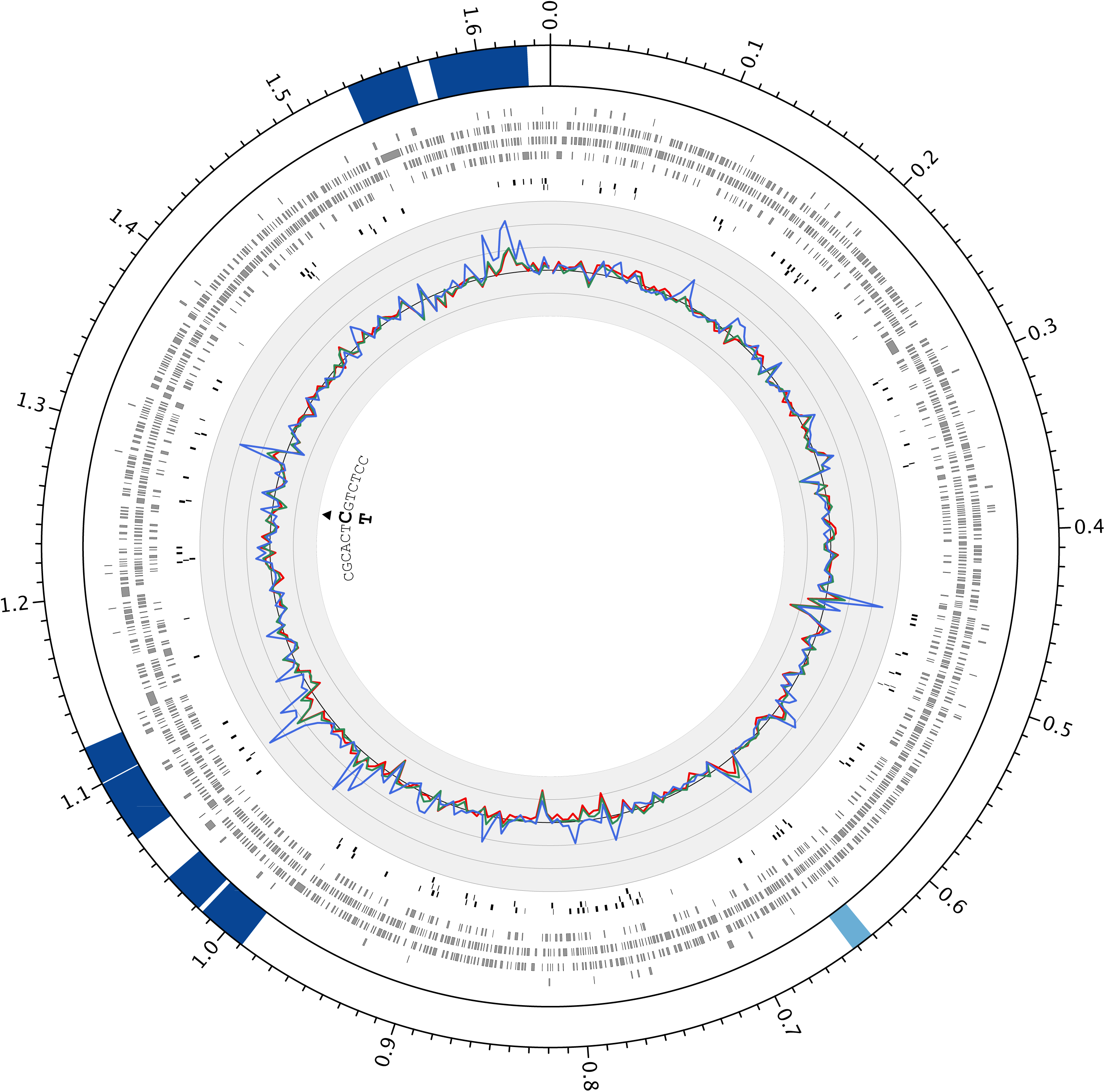
Circos plot of the *w*VulC *Wolbachia* genome. Prophage-derived regions are indicated inside the genome ideogram (outer track), with their colour corresponding to the score attributed by PHASTEST (dark blue: intact, light blue: questionable). Ticks on the ideogram are separated by 10 kb intervals, and indicated values are in Mb. Predicted protein-coding genes are represented by grey rectangles (second track), and Insertion Sequence elements by black rectangles (third track). In the fourth track, the normalized read coverage for 5 kb adjacent windows (averaged over recipient individuals) is represented for the different resequenced tissues: ovaries (red), nerve chain (green) and haemolymph (blue). The black line corresponds to a mean normalized coverage of one, and grey lines are separated by 0.1 unit. Black triangle indicates the position of the detected variant and the bold letter (C and T) correspond to the two alleles.

### 3.5 Whole-genome resequencing and variant calling reveal *Wolbachia* genome stability

The variant calling performed using the whole-genome resequencing of tissues from three females naturally infected by *Wolbachia* (i.e., source lineage) detected 641 SNPs and short indels. However, 584 of these apparent variants were also detected in the resequenced genome of several *A. vulgare* lineages not infected by *Wolbachia* (validation procedure, step 1), meaning that these variants probably amounted to *Wolbachia* sequences inserted into the host nuclear genome. The presence of reads sequenced from uninfected hosts within a 300-bp window around the 57 remaining polymorphic regions resulted in the exclusion of 26 variants (validation procedure step 2). Of the remaining 31 SNPs and small indels, 7 were excluded because their regions were PCR-amplified in males not infected with *Wolbachia* (infection status confirmed by qPCR, validation procedure step 3). Finally, in the last step of variant validation, we excluded 23 variants due to anomalies detected, such as a high proportion of clipped reads and/or a high number of mismatches between reads and reference. Overall, only 0.15% (one SNP) of the originally detected variants were finally validated. This unique SNP was observed in a single individual (F_1999). The frequency of this variant varied slightly among the three tissues (**Table 1**). The mutation affected a protein-coding gene, with a non-synonymous substitution (**Table 1**, **Fig. 5**). The putative protein did not have any recognized functional domain. The variant was not detected in the F_1999 individual’s sisters, and was also absent from all females tested in the maternal line (traced back 6 generations, see appendix 1).

**Table 1.** Nucleotide difference between *Wolbachia* from three tissues of naturally infected *Armadillidium vulgare*. HAEM: haemolymph, NC: nerve chain, OVA: ovaries. Nucl: Nucleotide, AA: amino acid; Arg: arginine, Gln: glutamine

| Genome position | Individual | Tissue | Frequency | Type of change | Nucl. change | AA change | ORF effect | Gene name | Functional domains |
| --- | --- | --- | --- | --- | --- | --- | --- | --- | --- |
| 1269492 | F1_2015 | NC | 0 | Substitution | C -> T | Arg340Gln | Missense variant | ABOJ95_001275 | No recognized functional domain |
|  |  | OVA | 0 |  |  |  |  |  |  |
|  | F2_2015 | NC | 0 |  |  |  |  |  |  |
|  |  | OVA | 0 |  |  |  |  |  |  |
|  | F_1999 | HAEM | 0.466 |  |  |  |  |  |  |
|  |  | NC | 0.584 |  |  |  |  |  |  |
|  |  | OVA | 0.239 |  |  |  |  |  |  |

## 4. Discussion

Experimental trans-infections enabled us to study the influence of the tissue origin of *Wolbachia* on the early infection dynamics of the endosymbiont in different recipient host tissues, and the ability of each bacterial sub-population to establish a chronic infection. At the same time, regular monitoring of injected individuals over a period of six months shed light on the influence of *Wolbachia* on the survival rate of recipient hosts and did not reveal any difference between *Wolbachia* sub-populations with respect to their virulence. In addition, our results corroborate a previous report showing a similar survival rate between trans-infected and un-infected animals (Le Clec’h et al., 2012). As the source and recipient lines belonged to the same field-collected *A. vulgare* population, they should be ecologically similar for *Wolbachia*. Low virulence is therefore expected, as for most vertically transmitted endosymbionts transferred between closely related hosts (Le Clec’h et al., 2013).

Colonization of recipient female tissues was relatively fast, with a threefold increase in *Wolbachia* density between the second and sixth month after infection. At the end of the experiment, the bacterial load in the tissues of the recipient females was similar to, or even higher than, that observed in the tissues of naturally vertically infected females.

Nevertheless, although rapid, the dynamics of *Wolbachia* infection varied according to the tissue origin of the bacteria. Bacteria from the nerve chain colonized recipient hosts much less rapidly than bacteria from ovaries and haemolymph. Two months after infection with a similar dose of bacteria, *Wolbachia* load was more than twice as low in hosts injected with bacteria from nerve chain, compared with hosts infected by bacteria from the other two tissues. The growth rate of bacteria is known to largely vary according to the type and amount of nutrients available in their environment (Keiblinger et al., 2010; Liu et al., 2005). The lower growth rate of bacteria from nerve chain, compared with those from the other tissues, could therefore be explained by the fact that in naturally infected hosts, this tissue is less favourable to the development of *Wolbachia* (e.g., poorer in resources). If so, bacteria from the nerve chain could be subject to growth-limiting stress at the time of their horizontal transfer, which would explain their lower replication rate within the recipient host at the start of infection. However, the nerve chain was the recipient host tissue the most rapidly colonized by *Wolbachia* after trans-infection, and the one showing the highest bacterial load at each sampling point irrespective of the tissue origin of the bacteria. As well as showing no tropism towards the tissue of origin, this result suggests that nervous tissue is actually favourable to the development of *Wolbachia*.

Alternatively, the lower growth rate from bacteria originating from the nerve chain could be adaptive. Classic theory for the evolution of virulence is based on a trade-off between parasite growth, transmission and host survival, which predicts that higher growth increases not only transmission but also virulence (Alizon et al., 2009; Frank, 1996). *Wolbachia* transmission is essentially maternal (Turelli et al., 2018; Werren et al., 2008), but cases of non-maternal transmission have been observed *in natura* (Durand et al., 2024; Sanaei et al., 2021) and in the laboratory (Le Clec’h et al., 2013; Rigaud & Juchault, 1995). In a tissue not directly involved in parasite transmission, we predicted that virulence should constrain the parasite growth to low levels. Our results corroborate this prediction, as *Wolbachia* from the nerve chain of *A. vulgare* should not be transmissible – except maybe in rare cases of cannibalism and predation (Le Clec’h et al., 2013) – and may prove detrimental to the host if they proliferate. Uncontrolled colonization of nerve cells has indeed been shown to alter host behaviour in several *Wolbachia*-host pairs (Bi & Wang, 2020; Le Clec’h et al., 2012), and to lead to severe tissue degeneration and premature host death (Kosmidis et al., 2014; W. L. Le Clec’h et al., 2012; Min & Benzer, 1997; Strunov & Kiseleva, 2016). In tissues that allow parasite transmission, the balance between *Wolbachia* growth and host fitness should favour higher growth rates. In terrestrial isopods, both ovaries and haemolymph play a key role in both intra- and inter-host dissemination of *Wolbachia*. Ovaries are the route for vertical transmission and haemolymph, in addition to participating in the dissemination of the bacteria across host tissues (Braquart-Varnier et al., 2015; Sanaei et al., 2021), may also be involved in horizontal transfer via contact between injured individuals (Rigaud & Juchault, 1995).

Although initial differences were observed at two months post-infection, the tissue origin of the injected *Wolbachia* had no significant effect on bacterial load in the three tissues analysed four months later (i.e., 6 months post-injection). The convergence of growth rates over time suggests that the early phenotypic variability observed two months post-infection may result from phenotypic plasticity. This interpretation is further supported by the absence of recurrent small genomic variations associated with tissue types in naturally infected individuals from the source lineage. More broadly, SNP and small indel analyses revealed a single substitution detected across sequence data from seven tissue samples. This SNP, observed in one female (F_1999), was nevertheless present in all three *Wolbachia* subpopulations, with allelic frequencies ranging from 24% to 58% depending on the tissue. This mutation, located in a gene coding for a protein of unknown function, leads to an amino acid substitution. Although we currently have no information on the phenotypic impact of this mutation, amino acid substitution may have strong phenotypic effects in bacteria (e.g., (Abdelaal et al., 2009; Bacigalupe et al., 2019)).

The variant, although predominant in the nerve chain, was present at lower frequencies in the ovaries. Nevertheless, over a quarter of the *Wolbachia* population in this tissue carried the mutation. This suggests that some of the female’s oocytes may have been colonized by the variant and that, despite the suspicion of a strong bottleneck, it could have passed it on to its offspring (Chrostek & Teixeira, 2015). However, this hypothesis could not be verified in our study, as the female was sacrificed before reproducing. What is clear is that the absence of this SNP within the individual’s maternal lineage strongly suggests that this *Wolbachia* variant arose during its lifetime. The emergence of *Wolbachia* variants within just a few host generations or even during a host’s lifetime has been documented (Chrostek & Teixeira, 2018; Martinez & Sinkins, 2023; Namias et al., 2024; Newton & Sheehan, 2015). However, most reported variants were associated with structural genomic changes rather than point genomic mutations (e.g., (Chrostek & Teixeira, 2018; Namias et al., 2024)). We currently have no information on the presence of structural genomic changes within the different *Wolbachia* tissular subpopulations, but in agreement with reports from other symbiotic systems, the very low number of point mutations observed in our study tends to support the stability of the *Wolbachia* genome (Dainty et al., 2021; Huang et al., 2020; Ross et al., 2022; Trouche et al., 2024).

In conclusion, our study reveals that within-host environmental heterogeneity can lead to diverse phenotypes in the most widespread bacterial endosymbiont in animals, *Wolbachia*. This variability did not appear to result from genomic variation, but from phenotypic plasticity. From a methodological point of view, we showed that the detection of variants in endosymbiont populations requires considerable caution, as our conservative approach led us to exclude more than 99.85% of the initially called variants. We recommend the use of a rigorous step-by-step approach to eliminate spurious genetic variants caused by the presence of endosymbiont genomic sequences inserted into the host genome.

## Data accessibility statement

The R scripts and data supporting the conclusions of this article are available on the Figshare data repository (https://figshare.com/s/b1f84643b65b4f897c54). To ensure the stability and reliability of the command-lines used for genome assembly and SNP calling we have included all necessary information and options in the github repository (https://github.com/UMR-CNRS-7267/Wolbachia_endosymbiont_heterogeneity_paper). We have taken proactive measures to address potential volatility on GitHub by uploading a compressed archive of this GitHub methodology to the Figshare data repository. Raw sequencing data have been deposited in GenBank (*w*VulC genome assembly: BioProject PRJNA1116085; *Wolbachia* resequencing: PRJNA1117639).

## Author’s contributions

R.P. and R.C. conceived and designed the experiments. R.P., R.J., M.P. and M.R. performed the trans-infection experiments. RP, TU, WA performed the preliminary experiment. RP and C.D prepared the samples for resequencing and C.D, TB and TU carried out the PCRs. D.O. performed *Wolbachia* genome sequencing. Y.D. assembled the *Wolbachia* genome. R.P., Y.D., A. M. and J.P. analysed the data. R.P. wrote the first draft of the manuscript, and all authors contributed substantially to revision.

## Competing interests

We declare we have no competing interests.

## Funding

This work was funded by Agence Nationale de la Recherche Grants ANR-21-CE02-0004 (RESIST) to RC and ANR-20-CE02-004 (SymChroSex) to JP, and intramural funds from the CNRS and the University of Poitiers.

## Acknowledgements

We thank Isabelle Giraud for the extraction of the DNA used to sequence and assemble the genome of *Wolbachia w*VulC and Pierre Grève and Didier Bouchon for providing us with preliminary data so that we could fine-tune the bioinformatics analyses. We would like to thank Alexandra Lafitte for her invaluable work supervising the breeding of *A. vulgare*.

## Notes

https://figshare.com/s/b1f84643b65b4f897c54

https://github.com/UMR-CNRS-7267/Wolbachia_endosymbiont_heterogeneity_paper

### Competing Interest Statement

The authors have declared no competing interest.

### Summary of Updates

Here's a revised version of the manuscript that includes two new experiments. The first aims to confirm that the quantity of bacteria injected into the different recipient hosts is indeed similar, and the second to follow the appearance of the Wolbachia variant detected in the different bacterial subpopulations within the maternal line of one of the sequenced individuals.

