## Appendix 1 for "Tissue heterogeneity is associated with phenotypic but not genomic diversity in *Wolbachia* endosymbionts"

### Supplementary materials

(1) *Wolbachia* quantification in tissues of naturally infected (*A. vulgare*)
(2) Estimation of the proportion of live *Wolbachia* in filtered tissue solutions injected into recipient hosts.
(3) Primers used for the ARMS PCR & Results.

#### (1) *Wolbachia* quantification in tissues of naturally infected *A. vulgare*

Previous studies have shown that the density of *Wolbachia* in the tissues of naturally infected hosts was different between tissues (e.g., Le Clec'h et al 2017). Here, we aimed to inject similar quantities of bacteria in recipient hosts, independently of their tissue origin. To achieve this, it was necessary to adjust the concentration of bacteria per ng of DNA in the different tissue solutions before injection into recipient hosts. To calculate the dilution factors of the different solutions, we first estimated the *Wolbachia* density per ng of DNA in the nerve chain, ovaries and haemolymph of 15 females from the source line (WXw). Absolute quantification of *Wolbachia* was estimated by quantitative PCR as described in the "Materials and methods" section of the main text. The gene copy number of *wsp* was then estimated by calculation in reference to a standard curve. The total DNA quantity (i.e. host+*Wolbachia*) of each sample, measured by fluorescence-based Qubit quantitation assays (Invitrogen™ Qubit™ Fluorometer), was used to normalize *wsp* gene copy number. The results are thus given in number of *wsp* copies per ng of total DNA (see Le Clec'h et al 2012 doi.org/10.1371/journal.ppat.1002844).

*Wolbachia* load varied significantly between females' tissues (LRT = 35.06, P < 0.0001, **Figure S1**). *Wolbachia* was less abundant in the haemolymph than in the nerve chain and ovaries (LRT = 9.35, P = 0.002, LRT = 35.04, P < 0.0001, respectively). *Wolbachia* density was significantly lower in the nerve chain than in the ovaries (LRT = 15.57, P < 0.0001). On average, we found 1392 ± 206 bacteria per DNA ng in the haemolymph, 3781 ± 986 bacteria per DNA ng in the nerve chain and, 13897± 3548 bacteria per DNA ng in the ovaries. We therefore implemented a dilution procedure to standardize the concentration of *Wolbachia* in each tissue solution used to transfect the bacteria (see main text).

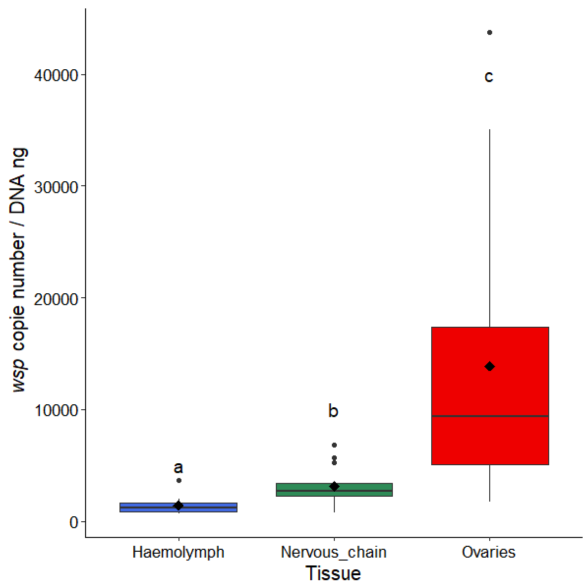


**Figure S1: *wsp* copy number per DNA ng in *Armadillidium vulgare* tissues**. Boxes above and below the medians (horizontal lines) show the first and third quartiles, respectively. Black diamonds represent the means. Levels not connected by the same letter are significantly different.

### (2) Estimation of the proportion of live *Wolbachia* in filtered tissue solutions injected into recipient hosts.

To estimate the proportion of live *Wolbachia* in filtered tissue solutions injected into recipient individuals, we initially opted for a flow cytometry approach with double labelling of bacteria with propidium iodide (PI, whose fluorescence is recorded in the ECD channel) and Syto24 (whose fluorescence is recorded in the FITC channel). This protocol works well for bacteria pure cultures or water samples (Berney et al., 2007, doi: 10.1128/AEM.02750-06). In our case, filtered tissue solutions constitute complex samples, (e.i., including numerous cell debris, mitochondria, bacteria), so it was impossible to identify a clearly defined bacterial population from the classical FSC vs SSC plot. To circumvent this problem, we used the plot FSC vs Syto 24 to define a normalization gate encompassing the main populations common to all samples, infected by *Wolbachia* or not (example for nervous chain samples are displayed on Fig S2.A). Then, a population present in only all infected samples was defined as the *Wolbachia* one (Fig S2 B). Finally, IP staining was evaluated within the Wolbachia gate for each sample (Fig S2.C). Signal was very low, so that we didn't consider counting of dead *Wolbachia* cells as being strong enough for statistical analyses, nevertheless it seemed that IP staining was not really different between infected samples and those which are not, thus suggesting that viability of the bacteria was very weakly compromised.

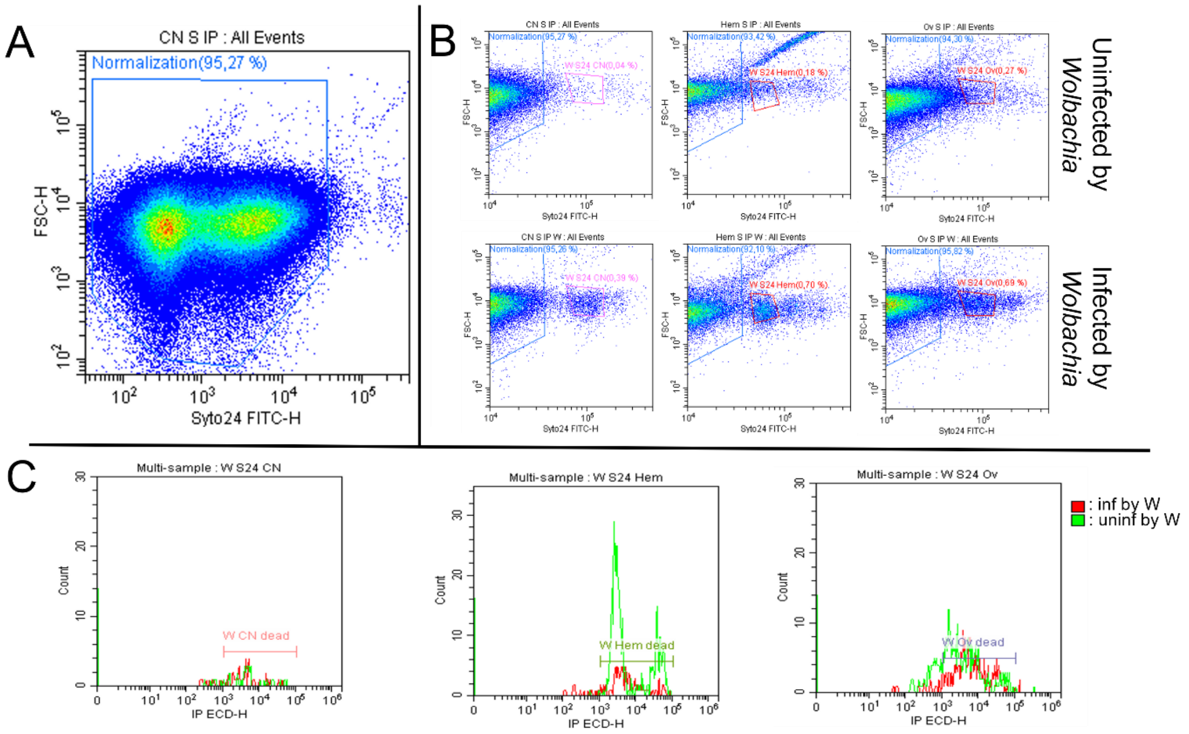


**Figure S2 Flow cytometry analysis of *Wolbachia*-infected and non-infected filtered tissue solutions** (**A**) set of events measured in a sample (here a filtered nerve chain solution). This figure illustrates the gating of the main populations use to normalize acquisition for every sample (245000 events counting within the Normalization gate). (**B**) Identification of the gate containing *Wolbachia*. To do this, we compared the scatterplots between tissue filtrates produced from infected females versus tissue filtrates produced from uninfected females. This figure illustrates the fact that the number of events that appear to be associated with *Wolbachia* is very low (less than 1% of total events). (**C**) Quantification of fluorescent events due to Propidium Iodide staining in the *Wolbachia* gate. (CN, nervous chain; Hem, hemocytes; Ov, ovary; W, *Wolbachia* infected). This entire experiment was replicated three times without ever giving better results.

We used an alternative molecular approach to infer bacterial viability indirectly through RNA quantification (see Matsuda et al. 2007, doi:10.1128/AEM.01224-06). Specifically, we analyzed tissue-origin effects as a proxy for bacterial viability by comparing the ratio between the reverse transcriptase qPCR (RT-qPCR) cycle threshold (CT) values to DNA-based qPCR CT values, both performed on RNA and DNA extracted from the same tissue filtrates. To achieve this, we prepared filtered tissue solutions from five females, as presented in the main text. DNA and RNA were then extracted from 80µL of each solution. DNA and RNA extraction was performed using standard protocols (DNA extraction: Qiagen DNeasy 96 Blood & Tissue kit, RNA extraction: Macherey-Nagel Nucleozol). DNA-based qPCR was performed as described in the main text, and RT-qPCR was carried out using the Luna® Universal One-Step RT-qPCR Kit. Each sample was replicated (technical replicate and a total of four biological replicates were performed (i.e., four filtered ovary solutions, four filtered nerve chain solutions and one filtered hemolymph solution). This experiment highlighted that the proportion of live bacteria in the different filtered tissue solutions seems to be similar (LRT = 4.0279, $p = 0.1335$, Ratio ± 95%CI, CT RNA - CT DNA ratio: hemolymph = 0.815 ± 0.017, nervous chain = 0.875 ± 0.014, ovaries = 0.862 ± 0.037).

### (3) Primers used for the ARMS PCR & Results.

**-Nucleotide sequence containing the SNP** (highlighted in yellow)
CTGGATTTCCATGGACCTTATTTACTCTACGCACTCGTCTCCTCTTTGGAGGAGCAGGTGGGCACACAGCTACC
TTTGGTTTTTCAGTTTTGTCTTCTTTTTTACTCACTTTCTCCTGTTTCTGTGTTTCATTATTTTTGTGGTTTATAACA
AATTTAGGTGAAGTGTCTACTAATTTTTCTACAGGTAAAAGAGCACGTTCTTGCTT

- **Primers used to target the reference sequence** (i.e., the *Wolbachia* lineage reported in all infected females)

Forward primer: TCCATGGACCTTATTTACTCTACGCACAC

Reverse primer: GCAAGAACGTGCTCTTTTACCT

- **Primers used to target the variant**

Forward primer : TAATAATAATCCATGGACCTTATTTACTCTACGCACAT

Reverse primer : GCAAGAACGTGCTCTTTTACCT

<u>ARMS PCR results:</u>

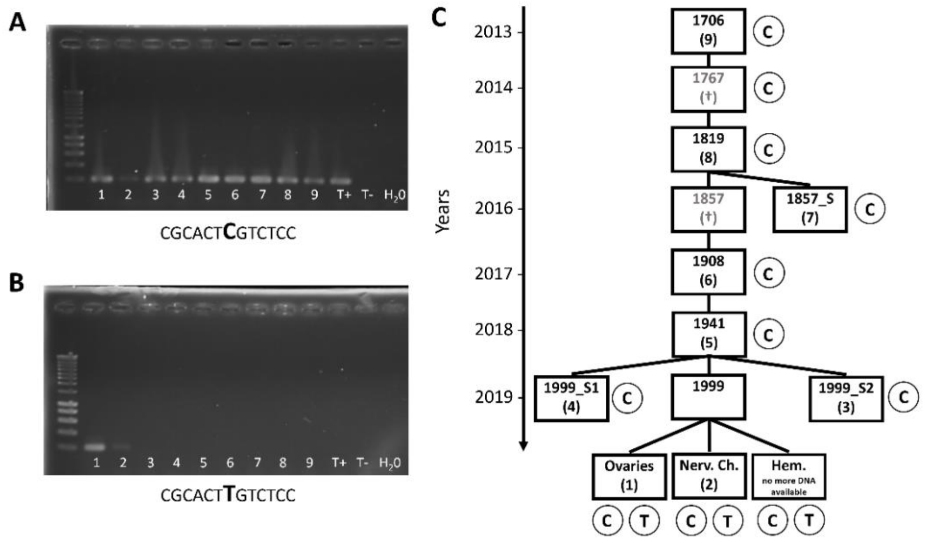


**Figure S3: ARMS PCR results and maternal genealogy of individual 1999.** (**A**) Agarose gel (1.5%) showing PCR products obtained using primers targeting the *Wolbachia* reference sequence. (**B**) Agarose gel (1.5%) showing PCR products obtained using primers targeting the *Wolbachia* variant sequence. Lanes 1–9 correspond to individual females, whose identities are shown in (**C**). **T+**: Control female (line WXw) infected with *Wolbachia* but unrelated to individual 1999. **T−**: Control female (line WXa) uninfected by *Wolbachia*. **$H_2O$**: Negative control. (**C**) Maternal genealogy of individual 1999, showing coinfection by two *Wolbachia* lineages as revealed by whole-genome resequencing on DNA from ovaries, nerve chain, and haemolymph (see table 1 in the main text). All the DNA extracted from the haemolymph of the 1999 female was used for whole-genome resequencing, which precluded validation of ARMS PCR in this sample. Two sisters of individual 1999 (1999_S1 and 1999_S2) were included in the analysis. If a direct maternal ancestor was unavailable (†: deceased before freezing), one of these sisters was used when possible. The circled letters represent the nucleotide observed at the variable position, with two letters indicating coinfection.
